# Multiple UBX proteins reduce the ubiquitin threshold of the mammalian p97-UFD1-NPL4 unfoldase

**DOI:** 10.1101/2022.01.13.476277

**Authors:** Ryo Fujisawa, Karim P.M. Labib

## Abstract

The unfolding of ubiquitylated proteins by the p97 / Cdc48 ATPase and its ubiquitin receptors Ufd1-Npl4 is essential in many areas of eukaryotic cell biology. Previous studies showed that yeast Cdc48-Ufd1-Npl4 is governed by a quality control mechanism, whereby substrates must be conjugated to at least five ubiquitins. Here we show that substrate processing by mammalian p97-UFD1-NPL4 involves a complex interplay between ubiquitin chain length and additional p97 cofactors. Using disassembly of the ubiquitylated CMG helicase as a model *in vitro* system, we find that reconstituted p97-UFD1-NPL4 only unfolds substrates with very long ubiquitin chains. However, this high ubiquitin threshold is greatly reduced, to a level resembling yeast Cdc48-Ufd1-Npl4, by the UBXN7, FAF1 or FAF2 partners of mammalian p97-UFD1-NPL4. Stimulation by UBXN7/FAF1/FAF2 requires the UBX domain that connects each factor to p97, together with the ubiquitin-binding UBA domain of UBXN7 and a previously uncharacterised coiled-coil domain in FAF1/FAF2. Furthermore, we show that deletion of the *UBXN7* and *FAF1* genes impairs CMG disassembly during S-phase and mitosis and sensitises cells to reduced ubiquitin ligase activity. These findings indicate that multiple UBX proteins are important for the efficient unfolding of ubiquitylated proteins by p97-UFD1-NPL4 in mammalian cells.

## Introduction

The regulated unfolding of proteins is essential for many aspects of cell biology and is mediated by multiple ATPases of the AAA+ protein family (Puchades, Sandate, & Lander, 2020; Seraphim & Houry, 2020; Zhang & Mao, 2020). These comprise hexameric rings with a central pore, in which loops from each subunit form a ‘spiral staircase’ around the unfolded polypeptide chain. ATP hydrolysis drives conformational changes within the ATPase hexamer, leading to translocation of the unfolded polypeptide along a ‘conveyor belt’ of pore loops.

p97 / Cdc48 / VCP is a AAA+ ATPase with orthologues in all three domains of life (N. Bodnar & T. Rapoport, 2017; van den Boom & Meyer, 2018). Eukaryotic p97 / Cdc48 / VCP unfolds highly stable proteins to prepare them for subsequent degradation by the proteasome (van den Boom & Meyer, 2018; Verma, Oania, Fang, Smith, & Deshaies, 2011; F. Wang, Li, Houerbi, & Chou, 2021). In addition, eukaryotic p97 / Cdc48 / VCP plays an essential role in diverse areas of cell biology by disrupting highly stable protein complexes or extracting proteins from cellular membranes. For example, p97 / Cdc48 / VCP is required for disassembly of the replisome at the end of chromosome replication (Dewar, Low, Mann, Raschle, & Walter, 2017; Franz et al., 2011; Maric, Maculins, De Piccoli, & Labib, 2014; Moreno, Bailey, Campion, Herron, & Gambus, 2014; Sonneville et al., 2017; Villa et al., 2021), release of newly synthesised protein phosphatase I from an inhibitory protein complex (van den Boom et al., 2021; Weith et al., 2018), release of DNA repair complexes from chromosomes (Kilgas et al., 2021; Meerang et al., 2011; van den Boom et al., 2016) and the proteasomal degradation of mis-folded membrane proteins (Ravanelli, den Brave, & Hoppe, 2020; Wolf & Stolz, 2012; Wu & Rapoport, 2018). Consistent with the important roles of p97 / Cdc48 / VCP in eukaryotic cell biology, dominant mutations in human p97 are associated with a late-onset multisystem proteinopathy (Johnson et al., 2010; Meyer & Weihl, 2014; Watts et al., 2004), whilst p97 is upregulated in certain cancer types (C. Li et al., 2021; Tsujimoto et al., 2004) and is a target for anti-cancer therapies (Anderson et al., 2015; Deshaies, 2014; Magnaghi et al., 2013; F. Wang et al., 2021).

p97 / Cdc48 / VCP associates with a wide range of adaptor proteins, many of which are still understood poorly (Buchberger, Schindelin, & Hanzelmann, 2015; Hanzelmann & Schindelin, 2017). Some recognise substrates directly (van den Boom et al., 2021; Weith et al., 2018), but others contain ubiquitin binding motifs that link p97 / Cdc48 / VCP to poly-ubiquitylated substrates (Bandau, Knebel, Gage, Wood, & Alexandru, 2012; Buchberger et al., 2015; Stach & Freemont, 2017). The best characterised ubiquitin adaptors of p97 / Cdc48 / VCP are Ufd1 and Npl4, which form a heterodimeric receptor for a specific class of polyubiquitin chains that are linked by lysine 48 (K48) of ubiquitin (N. Bodnar & T. Rapoport, 2017; Pan et al., 2021). Studies of yeast Cdc48-Ufd1-Npl4 have shown that Ufd1-Npl4 not only help to recruit Cdc48 to polyubiquitylated substrates but are directly involved in the first step of protein unfolding, which initiates within the ubiquitin chain rather than the substrate itself and is independent of ATP hydrolysis by Cdc48 (N. O. Bodnar & T. A. Rapoport, 2017; Ji et al., 2021; Twomey et al., 2019). Binding of Cdc48-Ufd1-Npl4 to the K48-linked chain induces unfolding of a ubiquitin moiety, with the unfolded ubiquitin polypeptide passing over a groove on the surface of Npl4 and thus being guided into the central pore of Cdc48 (Twomey et al., 2019). Subsequently, the pore loops drive ATP-dependent translocation, firstly of ubiquitin and then the attached substrate. In this way, Ufd1-Npl4 underpin a potentially universal mechanism for the recognition and unfolding of those Cdc48 substrates that are conjugated to K48-linked ubiquitin chains, without any requirement for direct recognition of the folded substrate protein itself (Ji et al., 2021; Twomey et al., 2019).

Structural studies indicated that the yeast Ufd1-Npl4 complex can bind simultaneously to four or five ubiquitins within the K48-linked chain on the substrate (Sato et al., 2019; Twomey et al., 2019). The functional significance of polyubiquitin binding was demonstrated by *in vitro* assays for disassembly of the 11-subunit CMG helicase (Deegan, Mukherjee, Fujisawa, Polo Rivera, & Labib, 2020), in which displacement of the ubiquitylated CMG subunit provided a readout for the number of conjugated ubiquitins that are required for substrate unfolding. Such assays showed that yeast Cdc48-Ufd1-Npl4 can only process substrates with at least five conjugated ubiquitins (Deegan et al., 2020). This ‘five-ubiquitin threshold’ likely represents a quality control mechanism that protects substrates from premature unfolding, by coupling Cdc48 activity to efficient ubiquitylation. Similarly, the recognition and degradation of ubiquitylated proteins by the proteasome is also governed by an analogous poly-ubiquitin threshold (Swatek & Komander, 2016).

Although Ufd1-Npl4 are essential for yeast Cdc48 and metazoan p97 / VCP to unfold ubiquitylated substrates (Blythe, Olson, Chau, & Deshaies, 2017; N. O. Bodnar & T. A. Rapoport, 2017; Deegan et al., 2020; Mukherjee & Labib, 2019), eukaryotic cells also encode a range of other adaptor proteins that combine ubiquitin-binding motifs with a UBX domain that associates with the amino terminus of p97 / Cdc48 / VCP (Alexandru et al., 2008; Hanzelmann & Schindelin, 2017; Kloppsteck, Ewens, Forster, Zhang, & Freemont, 2012; van den Boom & Meyer, 2018). Strikingly, studies of such ubiquitin-binding adaptors of human p97 found that UBXN7/UBXD7, FAF1, FAF2/UBXD8 and UBXN1/SAKS1 all co-purified with UFD1-NPL4 in addition to p97 (Alexandru et al., 2008; Hanzelmann, Buchberger, & Schindelin, 2011; Lee et al., 2013; Raman et al., 2015). These findings suggested that the mechanism or regulation of mammalian p97-UFD1-NPL4 might have additional layers of complexity, compared to yeast Cdc48-Ufd1-Npl4.

Here we show that the segregase activity of mammalian p97-UFD1-NPL4 is controlled by a combination of ubiquitin chain length and multiple UBX proteins. In contrast to yeast Cdc48-Ufd1-Npl4, reconstituted human p97-UFD1-NPL4 only unfolds substrates that are conjugated to extremely long ubiquitin chains. However, association of p97-UFD1-NPL4 with the UBX proteins UBXN7, FAF1 or FAF2 reduces the ubiquitin threshold to a comparable level to yeast Cdc48-Ufd1-Npl4. In this way, such UBX proteins ensure that p97-UFD1-NPL4 can unfold ubiquitylated proteins with high efficiency in mammalian cells.

## Results

### Mammalian p97-UFD1-NPL4 has a very high ubiquitin threshold

To investigate how ubiquitin chain length controls the ability of mammalian p97-UFD1-NPL4 to unfold substrate proteins, we adapted a reconstituted assay for the ubiquitylation and disassembly of budding yeast CMG helicase (Deegan et al., 2020), using purified recombinant versions of human p97 and UFD-NPL4 (Figure 1A-B). Initially, we used ubiquitylation conditions in which very long K48-linked ubiquitin chains were conjugated to the CMG-Mcm7 subunit (Deegan et al., 2020). The ubiquitylated helicase was then bound to beads that were coated with antibodies to the Cdc45 subunit, before incubation with human p97 and UFD1-NPL4 in the presence of ATP. Unfolding of ubiquitylated Mcm7 led to its release into the supernatant, together with all the other CMG subunits except for Cdc45 that remained bound to the antibody-coated beads (Figure 1C, compare lanes 1-2 and 3-4). Helicase disassembly required UFD1-NPL4 (Figure 1C, lanes 5-6) and was dependent upon the ATP binding and hydrolysis activity of p97 (Figure 1D-E).

**Figure 1.**
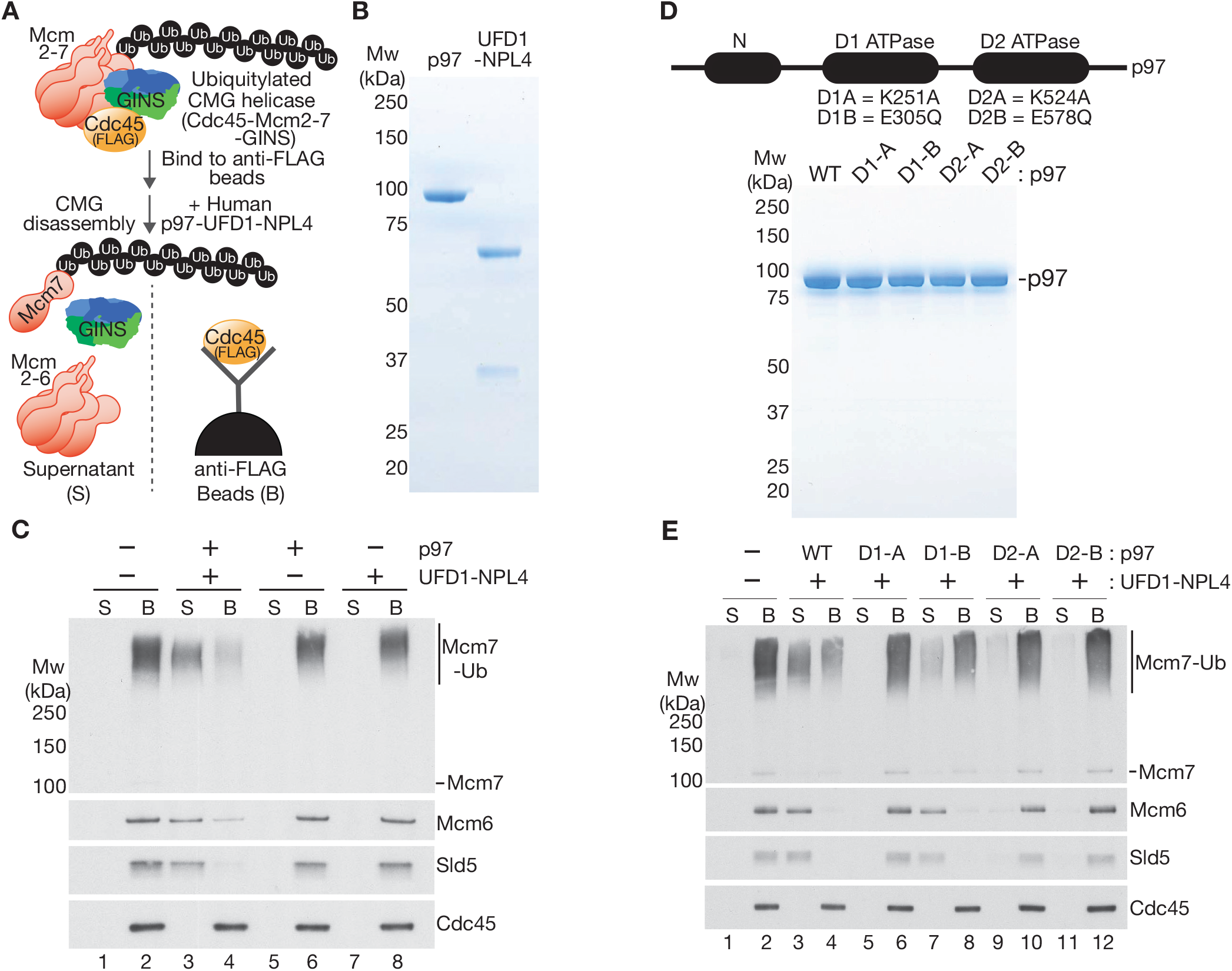
Reconstituted disassembly of ubiquitylated CMG helicase by purified human p97-UFD1-NPL4. (**A**) Budding yeast CMG helicase (with internal FLAG-tag on Cdc45 subunit) was ubiquitylated *in vitro*, bound to beads coated with anti-FLAG antibody and then incubated with recombinant human p97-UFD1-NPL4 (see Materials and Methods). Disassembly of ubiquitylated CMG displaced all subunits except Cdc45 into the supernatant. (**B**) Purified human p97 and UFD1-NPL4. (**C**) CMG disassembly reactions carried out according to the scheme in (A), in the presence of the indicated factors. (**D**) Purified recombinant human p97 with the indicated mutations in the D1 and D2 ATPase modules. (**E**) CMG disassembly reactions equivalent to those in (C), in the presence of the indicated factors.

Subsequently, we compared the ability of human p97-UFD1-NPL4 and yeast Cdc48-Ufd1-Npl4 to disassemble CMG complexes that either had very long ubiquitin chains attached to several lysine residues in CMG-Mcm7, or a single ubiquitin chain of up to ∼12 ubiquitins conjugated specifically to lysine 29 of CMG-Mcm7 (Figure 2A-B, see Materials and Methods). As described previously (Deegan et al., 2020), yeast Cdc48-Ufd1-Npl4 disassembled CMG complexes with five or more ubiquitins conjugated to CMG-Mcm7, though disassembly was most efficient when the attached ubiquitin chains were much longer (Figure 2C, compare lanes 3-4 and 5-6). Human p97-UFD1-NPL4 was comparably efficient to the yeast enzyme in disassembling CMG complexes with very long ubiquitin chains (Figure 2D, lanes 5-6). However, the human unfoldase was unable to disassemble CMG complexes that had up to ∼12 ubiquitins conjugated to CMG-Mcm7 (Figure 2D, lanes 3-4; also see Figure 3C, lanes 3-4). These findings indicated that human p97-UFD1-NPL4 has a much higher ubiquitin threshold than yeast Cdc48-Ufd1-Npl4.

**Figure 2.**
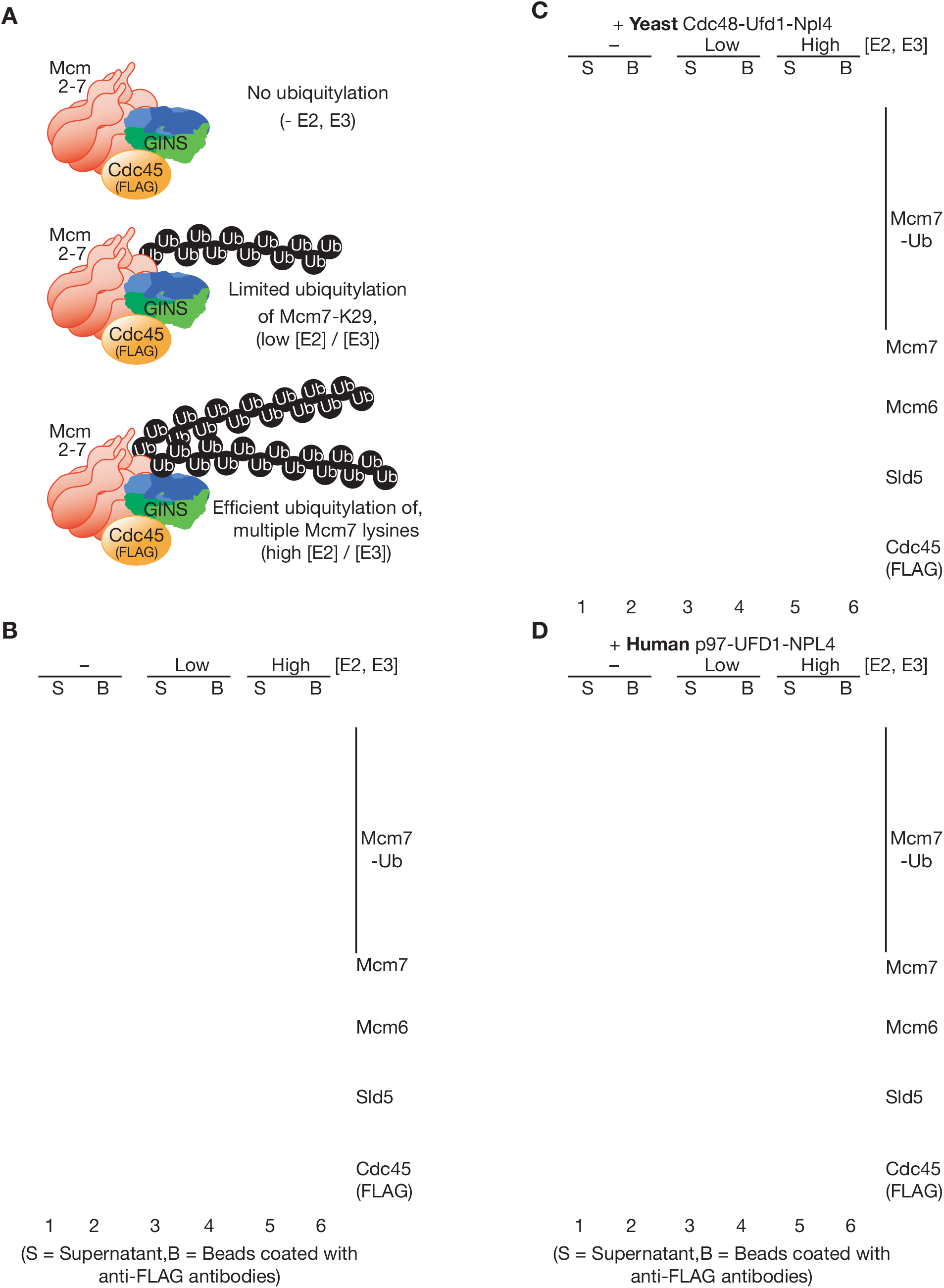
Human p97-UFD1-NPL4 has a much higher ubiquitin threshold than yeast Cdc48-Ufd1-Npl4. (**A**) Reconstituted CMG ubiquitylation reactions were performed under conditions that produced a single chain of up to ∼12 ubiquitins on lysine 29 of the CMG-Mcm7, or longer chains on multiple lysines of CMG-Mcm7 (see Materials and Methods). (**B**) The products of the ubiquitylation reactions in (A) were bound to beads coated with anti-FLAG antibodies. (**C**) Bead-bound ubiquitylated CMG from (B) was incubated with purified yeast Cdc48-Ufd1-Npl4. (**D**) Equivalent reactions in the presence of human p97-UFD1-NPL4.

**Figure 3.**
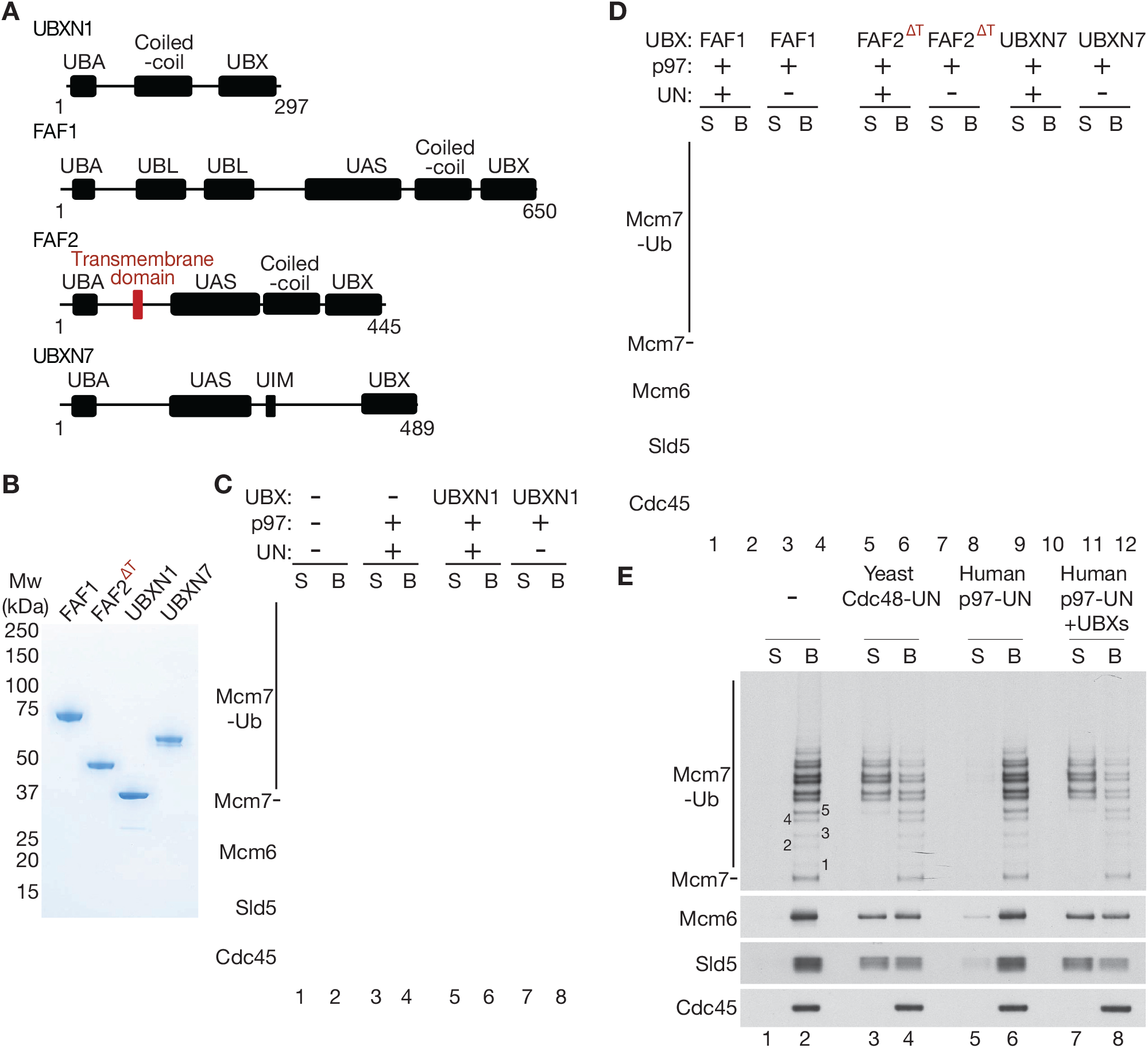
UBXN7, FAF1 and FAF2 reduce the ubiquitin threshold of p97-UFD1-NPL4. (**A**) Domain organisation of the indicated UBX proteins. UBA = ‘<u>UB</u>iquitin-<u>A</u>ssociated’ domain that binds ubiquitin; UBX = ‘<u>UB</u>iquitin regulatory <u>X</u>’ domain that binds p97; UBL = ‘UBiquitin-<u>L</u>ike’ domain that binds ubiquitin and NEDD8; UAS = domain of unknown function in FAF1 / FAF2 / UBXN7. (**B**) Purified proteins – the transmembrane domain of FAF2 was deleted in FAF2^ΔT^ to facilitate expression of a soluble protein. (**C**-**E)** CMG disassembly reactions in the presence of the indicated factors, performed as described above for Figures 1-2. See also Figure 3-figure supplements 1-2.

### UBXN7, FAF1 and FAF2 all reduce the ubiquitin threshold of mammalian p97-UFD1-NPL4

The UBX proteins FAF1, FAF2, UBXN7 and UBXN1 were previously found to co-purify with ubiquitin chains, p97 and UFD1-NPL4 from human cell extracts (Alexandru et al., 2008). Moreover, we found that all four factors bound directly to p97-UFD1-NPL4 *in vitro* (Figure 3-figure supplement 1; the association of each UBX protein with UFD1-NPL4 was dependent upon the presence of p97). Nevertheless, the interaction of UBXN1 with p97-UFD1-NPL4 was relatively weak, both *in vitro* with recombinant proteins (Figure 3-figure supplement 1) and in human cells (Alexandru et al., 2008).

We then monitored the impact of recombinant UBXN1, FAF1, FAF2 and UBXN7 on reconstituted CMG disassembly assays, under conditions where most CMG complexes had up to ∼12 ubiquitins conjugated to the Mcm7 subunit (Figure 3C-D). UBXN1 did not significantly alter the *in vitro* activity of the p97-UFD1-NPL4 unfoldase (Figure 3C), consistent with the weak interaction of UBXN1 with p97-UFD1-NPL4. In contrast, FAF1, FAF2 and UBXN7 all stimulated the disassembly of ubiquitylated CMG by human p97 (Figure 3D, lanes 1-2, 5-6, 9-10). In each case, disassembly was still dependent upon the presence of UFD1-NPL4 (Figure 3D, lanes 3-4, 7-8, 11-12). These data indicated that the UBX proteins FAF1, FAF2 and UBXN7 all reduce the ubiquitin threshold of mammalian p97-UFD1-NPL4 and thereby promote the efficient unfolding of ubiquitylated proteins. Notably, helicase disassembly in the presence of the three UBX proteins still required five or more ubiquitins on the CMG-Mcm7 subunit (Figure 3E), as seen previously for yeast Cdc48-Ufd1-Npl4 (Deegan et al., 2020).

### The UBX, UBA and UIM domains of UBXN7 all contribute to stimulation of p97-UFD1-NPL4 activity

To explore how UBXN7 stimulates the unfolding of ubiquitylated CMG-Mcm7 by p97-UFD1-NPL4, we purified a series of truncated versions (Figure 4A-B) and tested their ability to support CMG disassembly (Figure 4C). The UBX domain of UBXN7, which binds to the amino terminus of p97 (Z. H. Li, Wang, Xu, & Jiang, 2017), was essential for p97-UFD1-NPL4 to disassemble ubiquitylated CMG (Figure 4C, compare lanes 1-4). Moreover, the ubiquitin-binding UBA domain and the UIM domain that binds to ubiquitin and NEDD8 (den Besten, Verma, Kleiger, Oania, & Deshaies, 2012; Fisher et al., 2003; Young, Deveraux, Beal, Pickart, & Rechsteiner, 1998) both contributed to efficient CMG disassembly (Figure 4C, lanes 5-10). These findings indicated that UBXN7 binds to both p97-UFD1-NPL4 and the ubiquitin chain on CMG-Mcm7, thereby stabilising productive interactions between the unfoldase and the ubiquitylated substrate.

**Figure 4.**
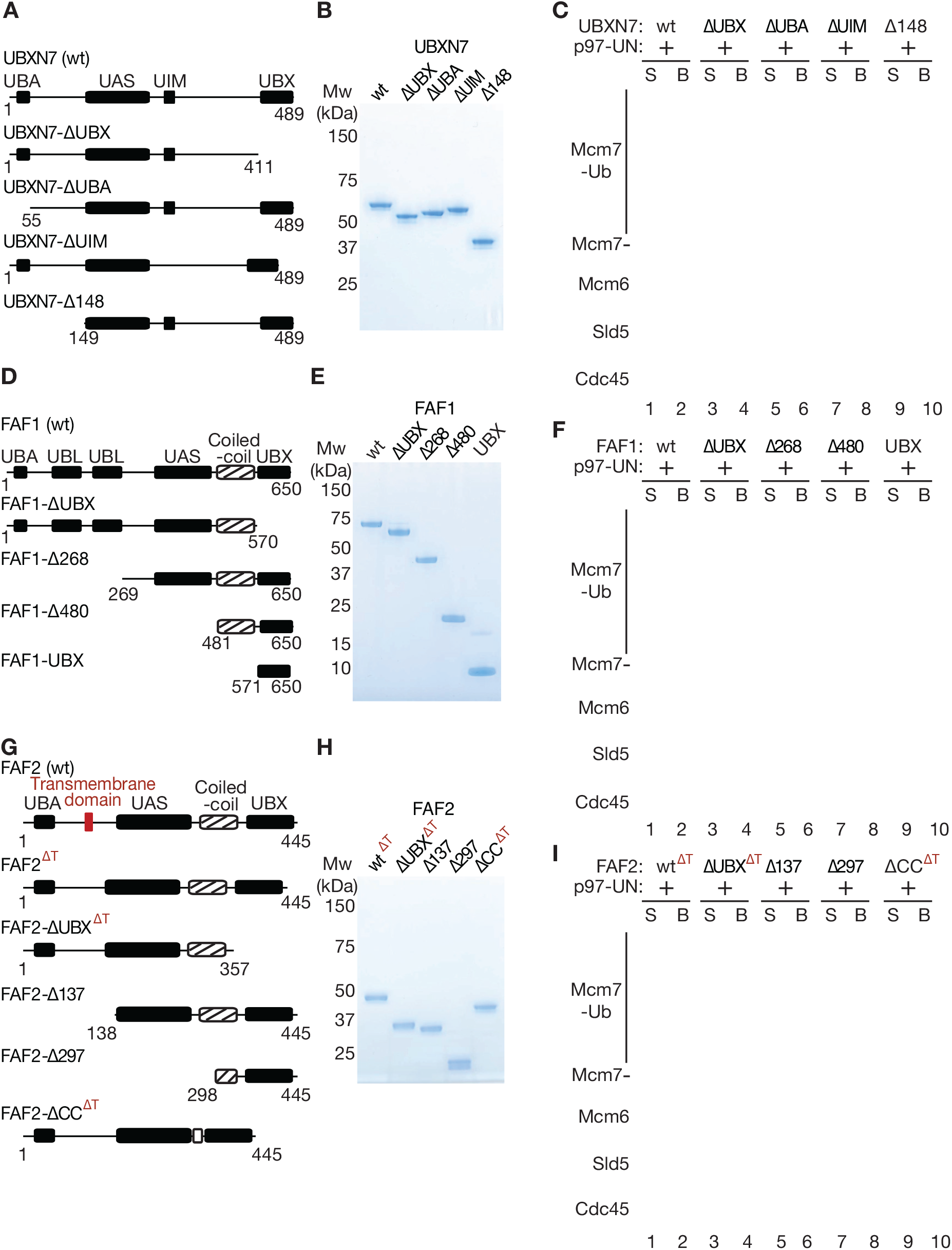
Mapping domains of UBXN7, FAF1 and FAF2 that stimulate the unfoldase activity of p97-UFD1-NPL4. (**A**) Truncations of UBXN7. (**B**) Purified proteins. (**C**) CMG disassembly reactions in the presence of the indicated factors, performed as described above. (**D**-**F**) Equivalent analysis for FAF1. (**G**-**I**) Analogous truncations of FAF2 – ‘ΔT’ indicates alleles that contain the amino terminus of the protein but lack the transmembrane domain. For (A), (D) and (G), the numbers correspond to residues in the full-length proteins. Domains were predicted using the SMART algorithm (http://smart.embl-heidelberg.de/) and Alphafold (Jumper et al., 2021).

### The UBX and coiled-coil domains of FAF1 and FAF2 are required to stimulate p97-UFD1-NPL4 unfoldase activity

An equivalent analysis of FAF1 truncations (Figure 4D-E) also showed that the UBX domain was essential to stimulate segregase activity (Figure 4F, compare lanes 1-4), indicating that FAF1 functions in a complex with p97-UFD1-NPL4, similarly to UBXN7. Surprisingly, however, the UBA domain of FAF1 was dispensable for stimulation of p97-UFD1-NPL4 (Figure 4F, FAF1-Δ268, lanes 5-6). Moreover, a carboxy-terminal fragment of FAF1 that comprised the coiled-coil and UBX domains was just as effective as the full-length protein in promoting disassembly of ubiquitylated CMG by human p97-UFD1-NPL4 (Figure 4F, FAF1-Δ480, lanes 7-8). In contrast, the UBX domain on its own was inactive (Figure 4F, FAF1-UBX, lanes 9-10). These findings indicated that the coiled-coil domain of FAF1 has a previously unanticipated role in stimulating the unfoldase activity of p97-UFD1-NPL4.

We also generated analogous truncations of FAF2 (Figure 4G-H) and again found that the UBX domain was essential for stimulation of the p97-UFD1-NPL4 unfoldase (Figure 4I, compare lanes 1-4), whereas the UBA domain was dispensable (Figure 4I, FAF2-Δ137, lanes 5-6). Furthermore, a FAF2 allele that lacked the coiled coil was inactive (Figure 4I, FAF2-ΔCC^ΔT^, lanes 9-10), whilst a fragment comprising the coiled-coil and UBX domains retained activity (Figure 4I, FAF2-Δ297, lanes 7-8). These findings indicated that FAF2 and FAF1 share a common mechanism by which they stimulate the disassembly of ubiquitylated CMG by p97-UFD1-NPL4.

Finally, we tested whether the novel role for the coiled-coil domain of mammalian FAF1 and FAF2 is also conserved in the single *C. elegans* orthologue UBXN-3. In reconstituted reactions analogous to those described above, we found that a carboxy-terminal fragment of UBXN-3 comprising the coiled-coil and UBX domains was sufficient to promote the disassembly of ubiquitylated CMG helicase by *C. elegans* CDC-48_UFD-1_NPL-4 (Figure 3-figure supplement 2A-B, UBXN-3-Δ435). Moreover, the UBX domain was essential for UBXN-3 to stimulate CDC-48_UFD-1_NPL-4 (Figure 3-figure supplement 2A-B, UBXN-3-ΔUBX), as reported recently (Xia, Fujisawa, Deegan, Sonneville, & Labib, 2021), but was insufficient in the absence of the coiled-coil domain (Figure 3-figure supplement 2A-B, UBXN-3-Δ527). Therefore, the coiled-coil domain is essential for UBXN-3 to stimulate *C. elegans* CDC-48_UFD-1_NPL-4, mirroring the importance of the coiled-coil domains of mammalian FAF1 and FAF2. These findings indicate that metazoan FAF1 / FAF2 / UBXN-3 stimulate p97-UFD1-NPL4 activity by a common mechanism, involving direct binding to the unfoldase and an essential role for the coiled-coil domain of the UBX cofactor.

### Cells lacking FAF1 and UBXN7 show defects in CMG helicase disassembly and are sensitive to inhibition of cullin ligase activity

Our data show that human p97-UFD1-NPL4 has a very high ubiquitin threshold *in vitro*, below which the unfoldase is dependent upon one of multiple UBX proteins, including UBXN7, FAF1 and FAF2. Such factors might act redundantly in mammalian cells, or else might stimulate p97-UFD1-NPL4 activity in distinct sub-cellular localisations. Previous work indicated that UBXN7 is largely nuclear in human cells (Raman et al., 2015). In contrast, FAF1 is present in both nucleus and cytoplasm (Raman et al., 2015), whereas FAF2 is a trans-membrane protein that localises to the endoplasmic reticulum, lipid droplets and the nuclear outer membrane (Go et al., 2021; Raman et al., 2015; Zehmer et al., 2009).

The function of human p97 and its cofactors is likely to be very highly conserved in other mammalian species, and we note that the mouse orthologues of p97-UFD1-NPL4, UBXN7, FAF1 and FAF2 are 97% - 100% identical to their human equivalents (Figure 5-figure supplements 1-2). To begin to assay for functional redundancy amongst the set of UBX proteins that stimulate mammalian p97-UFD1-NPL4 activity, we took advantage of the fact that mouse embryonic stem cells (mouse ES cells) were recently established (Villa et al., 2021) as a model system for studying mammalian CMG helicase disassembly. As in other metazoan species, mouse CMG is ubiquitylated on its MCM7 subunit during DNA replication termination, by the cullin ubiquitin ligase CUL2^LRR1^ (Villa et al., 2021). Ubiquitylated CMG is then disassembled very rapidly by p97-UFD1-NPL4, but the ubiquitylated helicase can be stabilised by treating cells with a small molecule inhibitor of the p97 ATPase (Villa et al., 2021).

CRISPR-Cas9 was used to make deletions in exon 1 of the *UBXN7* and *FAF1* genes (Figure 5-figure supplement 3), using mouse ES cells in which both copies of the *SLD5* gene had previously been modified to incorporate a GFP tag (Villa et al., 2021). The CMG helicase was then isolated from cell extracts, by immunoprecipitation of the GFP-tagged SLD5 subunit. In extracts of control or *FAF1Δ* cells, CMG-MCM7 ubiquitylation was scarcely detectable (Figure 5A, lanes 5-6). However, ubiquitylated CMG-MCM7 was observed in extracts of *UBXN7Δ* cells (Figure 5A, lane 7) and accumulated slightly in cells that lacked both FAF1 and UBXN7 (Figure 5A, lane 8). These data indicated that nuclear UBXN7 contributes to the disassembly of ubiquitylated CMG helicase during DNA replication termination in mouse ES cells, with a minor contribution from FAF1. Nevertheless, ubiquitylated CMG further accumulated when control or *UBXN7Δ FAF1Δ* cells were treated with p97 inhibitor (Figure 5B, lanes 5-8), suggesting that CMG disassembly in the absence of UBXN7 and FAF1 was not blocked completely.

**Figure 5.**
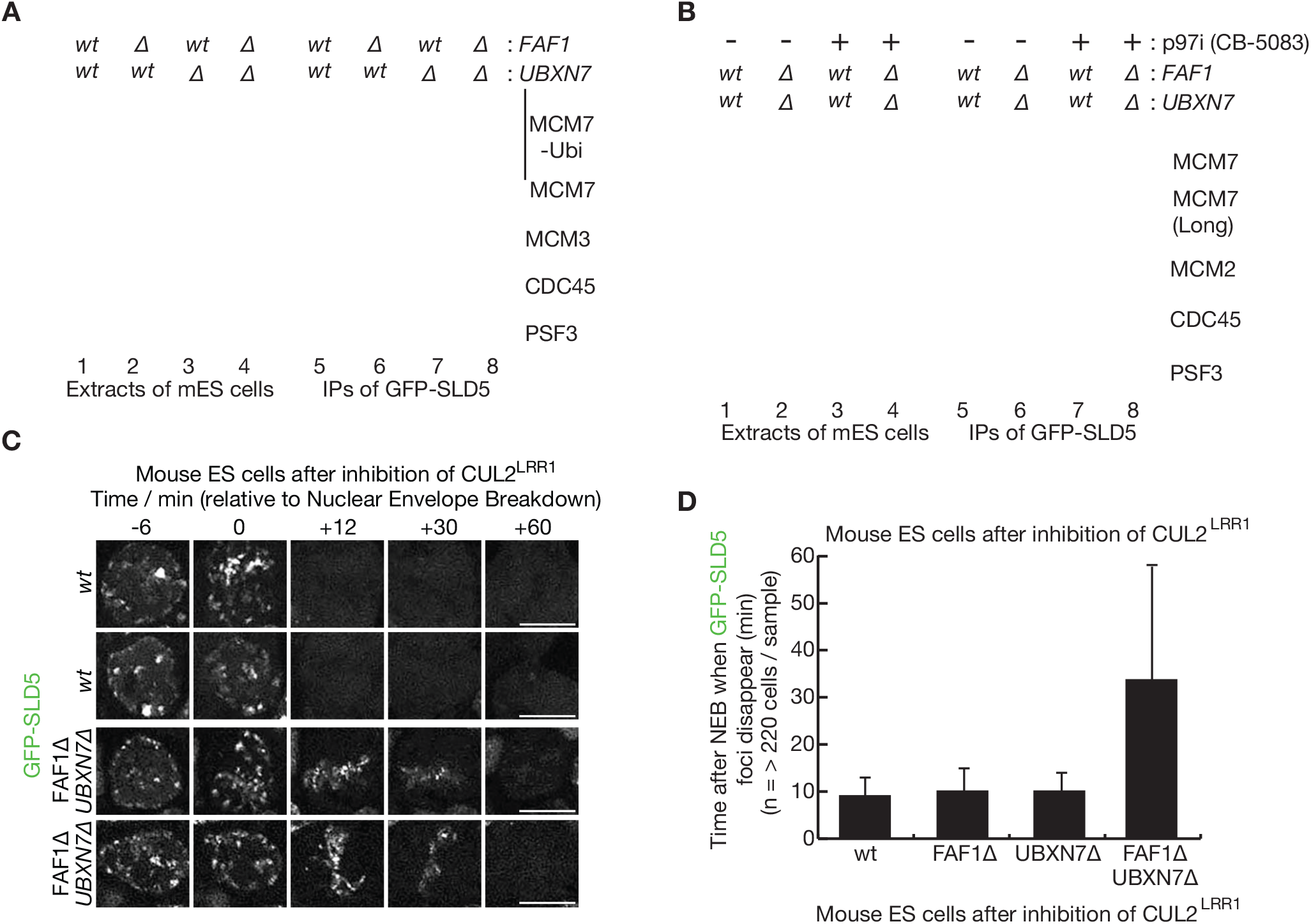
Partial redundancy between UBXN7 and FAF1 for CMG helicase disassembly during S-phase and mitosis in mouse ES cells. (**A**) CMG was isolated from extracts of mouse embryonic stem cells (mES) with the indicated genotypes, by immunoprecipitation of GFP-tagged SLD5 subunit of the helicase. (**B**) Equivalent experiment in which cells were treated as indicated with 5 µM CB-5083 (p97i = p97 inhibitor) for 3 hours before harvesting. (**C**) Time-lapse video analysis of mitotic entry in mouse ES cells expressing GFP-SLD5, following inhibition of CUL2^LRR1^. Scalebars correspond to 10 µm. (**D**) Quantification of the data in (C). The data represent the mean values for three biological replicates (experiments performed on different days), together with the corresponding standard deviations. See also Figure 5-figure supplements 1-4.

A second pathway for the disassembly of ubiquitylated CMG helicase acts during mitosis in metazoan cells and is important to process sites of incomplete DNA replication (Deng et al., 2019; Sonneville et al., 2019; Sonneville et al., 2017; Villa et al., 2021). This pathway is independent of CUL2^LRR1^ and instead requires the TRAIP ubiquitin ligase (Deng et al., 2019; Sonneville et al., 2019; Sonneville et al., 2017; Villa et al., 2021), which is mutated in human patients with a form of primordial dwarfism syndrome (Harley et al., 2016). To test whether FAF1 and UBXN7 also contribute to the mitotic disassembly of ubiquitylated CMG, we inactivated CUL2^LRR1^ in mouse ES cells and monitored the presence on mitotic chromatin of GFP-SLD5.

Time-lapse video analysis showed that CMG disassembly upon mitotic entry was blocked in the absence of TRAIP (Figure 5-figure supplement 4) as shown previously (Villa et al., 2021) and was about three times slower in *FAF1Δ UBXN7Δ* compared to *FAF1Δ*, *UBXN7Δ* or control cells (Figure 5C-D). Therefore, TRAIP-dependent CMG disassembly by p97 was delayed in cells that lacked both FAF1 and UBXN7, which act redundantly in the mitotic CMG disassembly pathway in mouse ES cells.

These findings suggested that the unfolding of other ubiquitylated substrates of p97-UFD1-NPL4 should also be impaired in the absence of FAF1 and UBXN7, particularly when ubiquitin chain formation on substrates of p97-UFD1-NPL4 is suboptimal. Cullin ligases represent the largest family of E3 enzymes directing K48-linked ubiquitin chain formation in eukaryotic cells (Harper & Schulman, 2021) and their activity is dependent upon neddylation of the cullin scaffold. Correspondingly, we found that *FAF1Δ UBXN7Δ* cells showed enhanced sensitivity to the neddylation inhibitor MLN4924 (Soucy et al., 2009), compared to *FAF1Δ*, *UBXN7Δ* or control cells (Figure 6). Overall, these findings suggest that the unfolding of ubiquitylated substrates by mammalian p97-UFD1-NPL4 is modulated by the length of the conjugated ubiquitin chain, together with the action of multiple UBX proteins.

**Figure 6.**
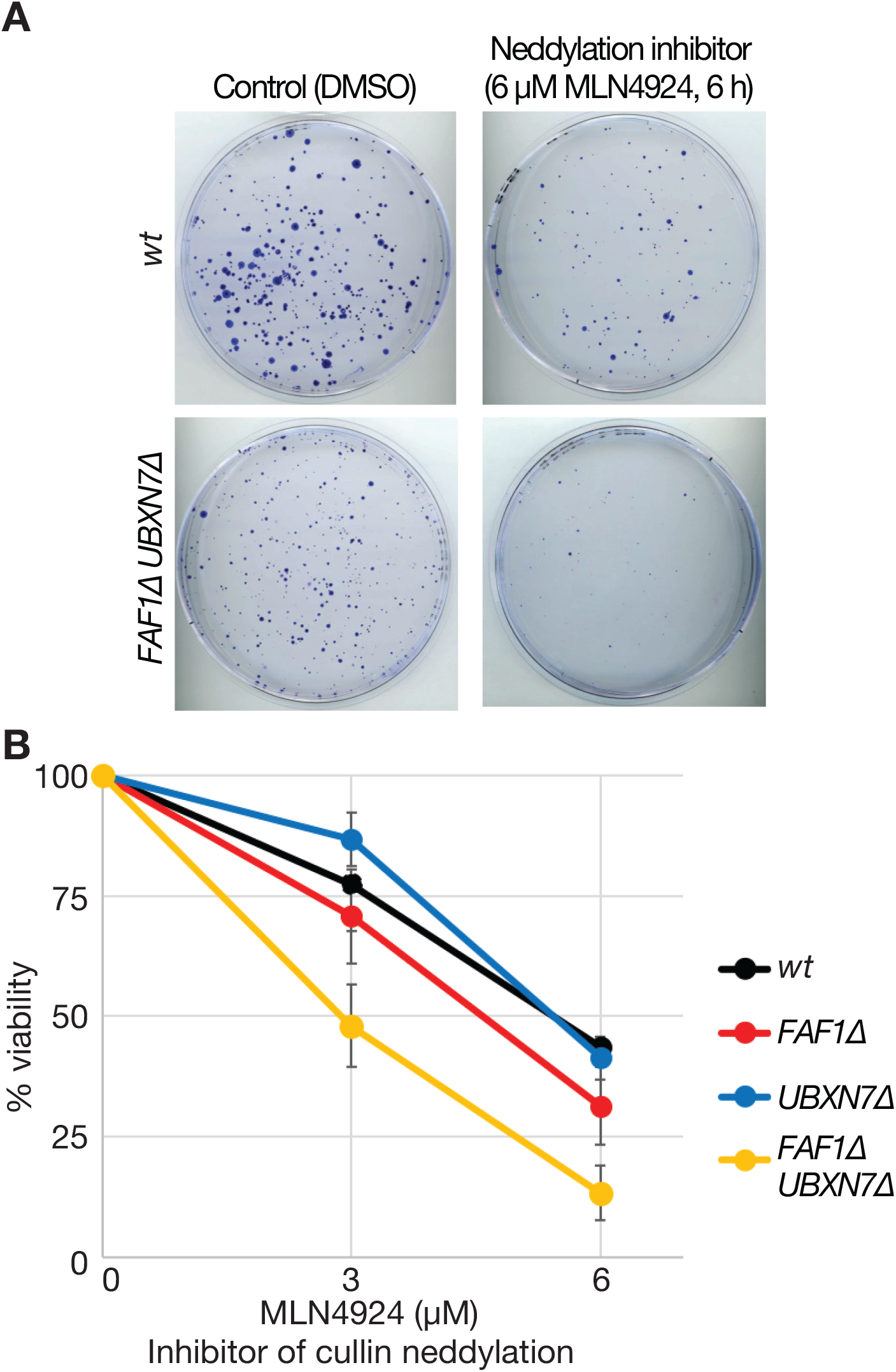
Cells lacking FAF1 and UBXN7 are sensitive to global inhibition of cullin ligase activity. (**A**) Cells of the indicated genotypes were treated as shown and then grown on 10 cm plates for 6 days. Surviving colonies were fixed and stained with crystal violet. (**B**) Viability of cells treated with 0, 3 or 6 µM MLN-4924 for six hours. The data represent the mean values and standard deviation for three biological replicates.

## Discussion

Our findings indicate that UBX proteins not only help to target p97 to substrates in a variety of sub-cellular locations, but also increase the efficiency of the p97-UFD1-NPL4 unfoldase and play a direct role in the unfolding reaction. UBXN7, FAF1 and FAF2 bind via their UBX domains to the amino terminal face of the p97 hexamer (Kim, Kang, Suh, & Yang, 2011; Z. H. Li et al., 2017), and by analogy with UFD1-NPL4 it is likely that the UBX proteins modulate substrate recognition and the initiation of substrate unfolding. Structural biology will play a central role in elucidating further the underlying mechanisms, but our data suggest that UBXN7 stabilises productive complexes between p97-UFD1-NPL4 and ubiquitylated substrates, with the UBXN7-UBX domain binding the unfoldase and the UBXN7-UBA domain binding to the ubiquitin chain. It is possible that FAF1 and FAF2 also assist in the recognition and initial unfolding of the substrate-linked ubiquitin chain by the p97-UFD1-NPL4 unfoldase. Nevertheless, the UBA domain of FAF1/FAF2 is dispensable for this function, which instead requires the coiled-coil domain of FAF1/FAF2, which is adjacent to the UBX domain and is functionally conserved in *C. elegans* UBXN-3. The mechanism by which the coiled coil of FAF1/FAF2 stimulates p97-UFD1-NPL4 remains to be explored in future studies, but one possibility is suggested by studies of the proteasomal AAA+ ATPase of Mycobacterium tuberculosis, which uses coiled coils to recognise the prokaryotic ubiquitin-like protein known as Pup (T. Wang, Darwin, & Li, 2010). By analogy, the coiled coil of FAF1/FAF2/UBXN-3 might represent a novel ubiquitin-binding domain that helps to stabilise an early intermediate in the unfolding of ubiquitylated substrates by p97-UFD1-NPL4.

In the presence of UBXN7, FAF1 and FAF2, p97-UFD1-NPL4 unfolds substrates with at least five conjugated ubiquitins (Figure 3), recapitulating the previously described ubiquitin threshold of yeast Cdc48-Ufd1-Npl4 (Deegan et al., 2020). This conserved five-ubiquitin threshold likely reflects the essential action of at least five ubiquitin binding modules within the UFD1-NPL4 heterodimer (Twomey et al., 2019), which together provide a form of quality control that prevents the premature or unscheduled unfolding of substrate proteins.

Our data indicate some degree of functional redundancy between UBXN7 and FAF1 in mammalian cells (Figures 5-6), which is likely to be mitigated by differences in their subcellular localisation (Raman et al., 2015), together with additional unique features of each factor such as the preferential association of the UBXN7-UIM domain with neddlyated cullins (den Besten et al., 2012). The growth of *UBXN7Δ FAF1Δ* cells is further impaired in the absence of FAF2 (Figure 5-figure supplement 3E), which likely regulates p97-UFD1-NPL4 on the surface of the endoplasmic reticulum, lipid droplets and the nuclear outer membrane (Go et al., 2021; Raman et al., 2015; Zehmer et al., 2009). The remaining activity of p97-UFD1-NPL4 in *UBXN7Δ FAF1Δ FAF2Δ* cells might involve additional UBX proteins that associate with p97-UFD1-NPL4 (Raman et al., 2015) and remain to be characterised *in vitro*. However, our data also indicate (Figure 1) that substrates with very long ubiquitin chains can be unfolded by p97-UFD1-NPL4 in the absence of UBX proteins, consistent with previous *in vitro* assays with an artificial model substrate (Blythe et al., 2017). Nevertheless, *UBXN7*, *FAF1* and *FAF2* all become essential for viability during mouse development or soon after birth (Adham et al., 2008; Koscielny et al., 2014), indicating their importance for the biology of mammalian p97-UFD1-NPL4. Similarly, the *ubxn-3* orthologue of mammalian FAF1 is important for embryonic viability and adult fertility during *C. elegans* development (Xia et al., 2021). Our data suggest that UBX proteins such as UBXN7, FAF1 and FAF2 all increase the efficiency of substrate unfolding by p97-UFD1-NPL4. We suggest that such factors are particularly important in mammalian cells whenever the length of the ubiquitin chain on the substrate is below the inherently high ubiquitin threshold of the mammalian unfoldase.

## Materials and Methods

Reagents and resources used in this study, including all purified proteins, are listed in Appendix Table S1.

### Plasmid DNA construction

The plasmids generated in this study are shown in Appendix Table S2. PCRs were conducted with Phusion® High-Fidelity DNA Polymerase (New England Biolabs, M0530), and amplified DNA fragments were cloned via the Gibson Assembly Cloning Kit (New England Biolabs, E2611) or with restriction enzymes and T4 ligase. All new constructs were verified by Sanger sequencing. Site-specific mutagenesis was performed using the Phusion polymerase, according to the manufacturer’s protocol.

Coding sequences of human p97 (Uniprot identifier P55072-1), UFD1 (Uniprot identifier Q92890-2), NPL4 (Uniprot identifier Q8TAT6-1), FAF1 (Uniprot identifier Q9UNN5-1) and UBXN7 (Uniprot identifier O94888-1) were amplified from XpressRef Universal Total human RNA (QIAGEN, 338112) by RT-PCR (TaKaRa, RR014) and cloned by Gibson assembly into the vector pK27SUMO (expressing a 14His-tagged version of the yeast SUMO protein Smt3) or pET28c. The coding sequences of human UBXN1 (Uniprot identifier Q04323-1) and FAF2 (Uniprot identifier Q96CS3-1) were synthesized by GenScript Biotech. For FAF2, the residues encoding the transmembrane domain (amino acids 90-118) were removed by PCR to improve solubility of the expressed protein.

### Expression of proteins in bacterial cells

Each plasmid was transformed into Rosetta (DE3) pLysS (Novagen, 70956), which was grown in LB medium supplemented with kanamycin (50 µg / ml) and chloramphenicol (35 µg / ml). Subsequently, a 100 ml culture was grown overnight at 37°C with shaking at 200 rpm. The following morning, the culture was diluted five-fold with 400 ml of LB medium supplemented with kanamycin (50 µg / ml) and chloramphenicol (35 µg / ml) and then left to grow at 37°C until an OD600 of 0.8 was reached. Protein expression was then induced overnight at 18°C in the presence of 1 mM IPTG. Cells were harvested by centrifugation for 15 minutes in a JLA-9.1000 rotor (Beckman) at 5,180 x g and the pellets were stored at –20 °C.

### Protease inhibitors

One tablet of ‘Roche cOmplete EDTA-free protease inhibitor Cocktail’ (Roche, 11873580001) was either added directly to 20 ml of protein purification buffer or else dissolved in 1 ml of water to make 20X stock.

### Buffers

Lysis buffer: 50 mM Tris-HCl (pH 8.0), 0.5 M NaCl, 5 mM Mg(OAc)_2_ and 0.5 mM TCEP

Gel filtration buffer: 20 mM Hepes-KOH (pH 7.8), 0.15 M NaCl, 5 mM Mg(OAc)_2_, 0.3 M sorbitol and 0.5 mM TCEP

Reaction buffer: 25 mM Hepes-KOH (pH 7.6), 10 mM Mg(OAc)_2_, 0.02%(w/v) IGEPAL CA-630, 0.1 mg/ml BSA, 1 mM DTT

Wash buffer: 100 mM Hepes-KOH (pH 7.9), 100 mM KOAc, 10 mM Mg(OAc)_2_, 2 mM EDTA, 0.1%(w/v) IGEPAL CA-630

Cell lysis buffer: 100 mM Hepes-KOH (pH 7.9), 100 mM KOAc, 10 mM Mg(OAc)_2_, 2 mM EDTA, 10 %(w/v) Glycerol, 0.1 % (w/v) Triton X-100, 2 mM sodium fluoride, 2 mM sodium β-glycerophosphate pentahydrate, 10 mM sodium pyrophosphate, 1 mM sodium orthovanadate, 1 µg/ml LR-microcystin, 1 mM DTT and Roche cOmplete EDTA-free protease inhibitor Cocktail.

### Purification of human p97 and its ATPase mutants

Bacterial pellets expressing 14His-Smt3-p97 from 500 ml cultures were resuspended in 20 ml Lysis buffer supplemented with 40 mM imidazole, 0.1 mM ATP, Roche cOmplete EDTA-free protease inhibitor Cocktail. The samples were lysed by incubation with 1 mg / ml lysozyme on ice for 30 minutes, followed by sonication for 90 seconds (15 seconds on, 30 seconds off) at 40% on a Branson Digital Sonifier, then clarified by centrifugation at 10,000 x g for 30 minutes in an SS-34 rotor (Sorvall).

The supernatant was mixed with 1 ml resin volume of Ni-NTA beads (Qiagen, 30210) which was pre-equilibrated with Lysis buffer supplemented with 40 mM imidazole and 0.1 mM ATP. After 1 hour incubation at 4°C, beads were recovered in a disposable gravity flow column and washed with 150 ml of Lysis buffer containing 40 mM imidazole and 0.1 mM ATP, followed by 10 ml of Lysis buffer containing 60 mM imidazole, 2 mM ATP and 10 mM MgOAc. 14His-Smt3-p97 was eluted with 5 ml of Lysis buffer containing 400 mM imidazole and 0.1 mM ATP. Ulp1 protease (10 µg / ml) was added to the eluate and incubated at 4°C overnight, to cleave 14His-Smt3 from p97. The sample was then loaded onto a 24 ml Superose 6 column equilibrated in Gel filtration buffer containing 0.1 mM ATP. Fractions containing hexameric p97 were pooled, concentrated with an Amicon Ultra-4 Centrifugal Filter Unit (Merck, UFC803024) according to the manufacture’s protocol, aliquoted and snap-frozen with liquid nitrogen.

### Purification of human UFD1-NPL4

Human untagged / 6His-NPL4 and 14His-Smt3-UFD1 were expressed individually in 750 ml and 250 ml cultures (at a 3:1 ratio of NPL4 to UFD1). The respective bacterial pellets were then resuspended in 15 ml and 5 ml of Lysis buffer containing 30 mM imidazole, Roche protease inhibitor tablets. The samples were then mixed and lysed by incubation with 1 mg / ml lysozyme on ice for 30 min, followed by sonication for 90 seconds (15 seconds on, 30 seconds off) at 40% on a Branson Digital Sonifier, before clarification by centrifugation at 10,000 x g for 30 minutes in an SS-34 rotor.

The supernatant was mixed with 1 ml resin volume of Ni-NTA beads which had been pre-equilibrated with Lysis buffer containing 30 mM imidazole. After 1 hour incubation at 4°C, beads were recovered in a disposable gravity flow column and washed with 150 ml of Lysis buffer containing 30 mM imidazole, followed by 10 ml of Lysis buffer containing 50 mM imidazole, 2 mM ATP and 10 mM MgOAc. The complex of 14His-Smt3-UFD1 with NPL4 was eluted with 5 ml of Lysis buffer containing 400 mM imidazole.

The yeast Ulp1 protease (10 µg/ml) was then added to cleave the 14His-Smt3 tag from UFD1, and the mixture was dialyzed into 500 ml of Lysis buffer containing 30 mM imidazole at 4 °C overnight. The dialyzed mixture was mixed with 0.5 ml Ni-NTA beads equilibrated with Lysis buffer containing 30 mM imidazole. After 15 minutes of rotation at 4 °C, the flow-through fraction was collected and concentrated with an Amicon Ultra-4 Centrifugal Filter Unit according to the manufacture’s protocol and then loaded onto a 24 ml Superdex 200 column equilibrated in Gel filtration buffer. Fractions containing UFD1-NPL4 were pooled, concentrated, aliquoted and snap-frozen with liquid nitrogen.

### Purification of metazoan UBX proteins

Bacterial pellets expressing 14His-Smt3 tagged UBX proteins from 500 ml cultures were resuspended in 20 ml of Lysis buffer containing 30 mM imidazole and Roche cOmplete EDTA-free protease inhibitor Cocktail. The samples were lysed by incubation with 1 mg / ml lysozyme on ice for 30 minutes, followed by sonication for 90 seconds (15 seconds on, 30 seconds off) at 40% on a Branson Digital Sonifier, before clarification by centrifugation at 10,000 x g for 30 minutes in an SS-34 rotor.

The supernatant was mixed with 1 ml resin volume of Ni-NTA beads which was pre-equilibrated with Lysis buffer containing 30 mM imidazole. After 1 hour incubation at 4°C, beads were recovered in a disposable gravity flow column and washed with 150 ml of Lysis buffer containing 30 mM imidazole, followed by 10 ml of Lysis buffer containing 50 mM imidazole, 2 mM ATP and 10 mM MgOAc. 14His_-_Smt3 tagged UBX proteins were eluted with 5 ml of Lysis buffer containing 400 mM imidazole. Ulp1 protease (10 µg/ml) was then added to cleave the 14His-Smt3 tag from UBX proteins during an overnight incubation. Subsequently, the samples were concentrated with an Amicon Ultra-4 Centrifugal Filter Unit according to the manufacture’s protocol and loaded onto a 24 ml Superdex 200 column in Gel filtration buffer. In the case of the fragments comprising the UBX domain of FAF1 and UBXN-3, a Superdex75 column was used instead of Superdex 200. The peak fractions containing each UBX protein were pooled, aliquoted and snap-frozen with liquid nitrogen.

### Purification of budding yeast proteins

The budding yeast proteins used in this study (replisome factors, ubiquitylation factors, and Cdc48 plus its Ufd1-Npl4 adaptors) are described in Table S1. The various factors were expressed using plasmids and strains described in Table S1 and were purified as described previously (Deegan et al., 2020).

### Purification of *C. elegans* proteins

The *C. elegans* proteins used in this study (replisome factors, ubiquitylation factors, neddylation factors and CDC-48 plus adaptors) are described in Table S1. The various factors were expressed using plasmids and strains described in Table S1 and were purified as described previously (Xia et al., 2021), except for UBXN-3-Δ435 and UBXN-3-Δ527 that are depicted in Figure 3-figure supplement 2 and were expressed as fusions to 14His-Smt3 in *E. coli* using the plasmids pRF056 and pRF057, as described above for other UBX proteins.

### Reconstituted ubiquitylation and disassembly of yeast CMG helicase

Yeast CMG was ubiquitylated in reconstituted *in vitro* reactions as described previously (Deegan et al., 2020). In summary, 15 nM yeast CMG was mixed with 30 nM Uba1, the indicated concentrations of the E2 Cdc34 and E3 SCF^Dia2^, 30 nM Ctf4 and 6 µM ubiquitin, in ‘Reaction buffer’ containing 100 mM KOAc and 2 mM ATP. To produce long ubiquitin chains on multiple lysines of CMG-Mcm7 (Figure 1 and Figure 2 ‘high [E2] / [E3]’), we used 25 nM of the E2 Cdc34 and 10 nM E3 SCF^Dia2^. To produce shorter chains (Figure 2 ‘low [E2] / [E3]’) that were restricted to lysine 29 of CMG-Mcm7 (Deegan et al., 2020), we used 2.5 nM of Cdc34 and 1 nM SCF^Dia2^. Subsequently, 2 nM Cdc34 and 2 nM SCF^Dia2^ were used for the experiments in Figure 3C and 3D, whereas 1.5 nM Cdc34 and 2 nM SCF^Dia2^ were used for the experiments in Figure 4C, Figure 4F and Figure 4I. Ubiquitylation reactions were conducted at 30°C for 20 minutes and then stopped by the addition of KOAc to a final concentration of 700 mM.

The ubiquitylated yeast CMG was then incubated for 30 minutes at 4°C with 2.5 µl (per 10 µl ubiquitylation reaction mix) magnetic beads (Dynabeads M-270 Epoxy; Life Technologies, 14302D) that had been coupled to anti-FLAG M2 antibodies. After the incubation, the bead-bound protein complexes were washed twice with 190 µl of Reaction buffer containing 100 mM KOAc and then resuspended in Reaction buffer containing 100 mM KOAc and 5 mM ATP. For the disassembly reactions, 50 nM each of human p97 (or yeast Cdc48), human UFD1-NPL4 (or yeast Ufd1-Npl4), and the indicated human UBX proteins were added to the suspension of beads bearing ubiquitylated yeast CMG, before incubation at 30°C for 20 minutes with shaking at 1000 rpm on an Eppendorf ThermoMixer®. The supernatants were collected and the beads washed twice with 190 µl of Reaction buffer containing 700 mM KOAc, before elution of bound proteins by the addition of LDS-PAGE sample loading buffer (Invitrogen, NP0007) and heating for 3 minutes at 95°C.

### Reconstituted ubiquitylation and disassembly of *C. elegans* CMG helicase

For the experiment in Figure 3-figure supplement 2B, worm CMG was ubiquitylated in reconstituted *in vitro* reactions as described previously (Xia et al., 2021), using proteins described in Appendix Table S1. In summary, 15 nM worm CMG was mixed with 50 nM UBA-1, 300 nM LET-70, 300 nM UBC-3, 50 nM ULA-1_RFL-1, 300 nM UBC-12, 100 nM DCN-1, 500 nM NED-8, 15 nM CUL-2^LRR-1^, 60 nM TIM-1_TIPIN-1, 30 nM POLε, 30 nM CLSP-1, 30 nM MCM-10, 60 nM CTF-18_RFC, 20 nM CTF-4 and 10 µM ubiquitin in ‘Reaction buffer’ containing 100 mM KOAc and 5 mM ATP. Ubiquitylation reactions were conducted at 20°C for 20 min.

The ubiquitylated worm CMG was then incubated for 30 minutes at 4°C with 2 µl (per 10 µl ubiquitylation reaction mix) magnetic beads (Dynabeads M-270 Epoxy; Life Technologies, 14302D) that had been coupled to anti-worm SLD-5 antibodies. After the incubation, the bead-bound protein complexes were washed twice with 190 µl of Reaction buffer containing 100 mM KOAc and then resuspended in Reaction buffer containing 5 mM ATP. For the disassembly reactions, 200 nM of *C. elegans* CDC-48.1, 50 nM of *C. elegans* UFD-1_NPL-4.1 and 50nM of the indicated form of *C. elegans* UBXN-3 were added to the suspension of beads bearing ubiquitylated worm CMG. The reactions were then incubated at 20°C for 20 minutes, whilst shaking at 1000 rpm on an Eppendorf ThermoMixer®. The supernatants were then collected and the beads washed twice with 190 µl of Reaction buffer containing 700 mM KOAc. The bound proteins were eluted by addition of LDS-PAGE sample loading buffer (Invitrogen, NP0007) and heating for 3 minutes at 95°C.

### Monitoring the interaction of UBX proteins with p97-UFD1-NPL4

For the experiment in Figure 3-figure supplement 1, 3 µg of 6His-NPL4_UFD1, 3 µg of each UBX protein and 13 µg of p97 were mixed as indicated with 2 µl slurry of Ni-NTA beads in a total of 11 µl using ‘Gel filtration buffer’ containing 30 mM imidazole and 2 mM ATP. After 15 minutes incubation on ice, the Ni-NTA beads were washed three times with 200 µl of ‘Wash buffer’ containing 20 mM imidazole and 0.5 mM ATP. Bead-bound proteins were eluted in LDS-PAGE sample loading buffer containing 50 mM EDTA, by heating for 3 minutes at 95°C.

### SDS-PAGE and Immunoblotting

Protein samples were resolved by SDS–polyacrylamide gel electrophoresis in NuPAGE Novex 4 -12% Bis-Tris gels (ThermoFisher Scientific, NP0321 and WG1402A) either with NuPAGE MOPS SDS buffer (ThermoFisher Scientific, NP0001) or NuPAGE MES SDS buffer (ThermoFisher Scientific, NP0002), or with NuPAGE Novex 3 - 8% Tris-Acetate gels (ThermoFisher Scientific, EA0375BOX and WG1602BOX) using NuPAGE Tris-Acetate SDS buffer (ThermoFisher Scientific, LA0041). Resolved proteins were either stained with InstantBlue (Expedeon, ISB1L) or were transferred to a nitrocellulose membrane with the iBlot2 Dry Transfer System (Invitrogen, IB21001S). Antibodies used for protein detection in this study are described in Appendix Table S1. Conjugates to horseradish peroxidase of anti-sheep IgG from donkey (Sigma-Aldrich, A3415), or anti-mouse IgG from goat (Sigma-Aldrich, A4416) were used as secondary antibodies before the detection of chemiluminescent signals on Hyperfilm ECL (cytiva, 28906837, 28906839) with ECL Western Blotting Detection Reagent (cytiva, RPN2124).

### Preparation of antibody-coated magnetic beads

Firstly, 300 mg of Dynabeads M-270 Epoxy (ThermoFisher Scientific, 14302D) were resuspended in 10 ml of dimethylformamide. Subsequently, 425 µl slurry of activated magnetic beads, which corresponded to ∼ 1.4 x 10^9^ beads, were washed twice with 1 ml of 1 M NaPO_3_ (pH 7.4). The beads were then incubated with 300 µl of 3 M (NH4)_2_SO_4_, 300 µg of anti-FLAG M2 antibody (Sigma-Aldrich, F3165), and 1 M NaPO_3_ (pH 7.4) up to a total volume of 900 µl. The mixture was then incubated at 4 °C for 2 days with rotation. Finally, the beads were treated as follows: four washes with 1 ml PBS; 10 minutes in 1 ml PBS containing 0.5 %(w/v) IGEPAL CA-630 with rotation at room temperature; 5 minutes in 1 ml PBS containing 5 mg / ml BSA with rotation at room temperature; one wash with 1 ml PBS containing 5 mg / ml BSA. The beads were then resuspended with 900 µl PBS containing 5 mg/ml BSA and stored at 4°C.

### Growth of mouse ES cells

Mouse ES cells were grown as described previously (Villa et al., 2021). E14tg2a cells (Appendix Table S1) were cultured in a humidified atmosphere of 5 % CO_2_, 95 % air at 37 °C under feeder-free conditions with Leukemia Inhibitory Factor (LIF; MRC PPU Reagents and Services DU1715) in serum-containing medium. All culturing dishes were precoated with PBS containing 0.1 % (w/w) gelatin (Sigma-Aldrich, G1890) for at least 5 minutes prior to seeding. The medium was based on Dulbecco’s Modified Eagle Medium (DMEM; ThermoFisher Scientific, 11960044), supplemented with 10 % Fetal Bovine Serum (FBS, FCS-SA/500, Labtech), 5 % Knockout serum replacement (ThermoFisher Scientific, 10828028), 2 mM L-Glutamine (ThermoFisher Scientific, 25030081), 100 U/ml Penicillin-Streptomycin (ThermoFisher Scientific, 15140122), 1 mM Sodium Pyruvate (ThermoFisher Scientific, 11360070), a mixture of seven non-essential amino acids (ThermoFisher Scientific, 11140050), 0.05 mM β-mercaptoethanol (Sigma-Aldrich, M6250) and 0.1 µg/ml LIF. For passaging, cells were released from dishes using 0.05 % Trypsin-EDTA (ThermoFisher Scientific, 25300054).

To determine doubling times, 200,000 or 500,000 cells (initial cell number N1) were grown for 48 hours on a 6-well plate, before recovery with 0.05 % Trypsin / EDTA. The total cell number in the final suspension (final cell number N2) was determined using a CellDrop BF (DeNovix). The doubling time (G) was then calculated via the formula G = (48 x log(2)) / (log(N2) – log(N1)). The experiments were performed in triplicate and mean values were determined, together with the standard deviation (SD).

### CRISPR-Cas9 genome editing

To design a pair of guide RNAs (gRNAs) to target a specific site in the mouse genome, we used ‘The CRISPR Finder’ provided by the Welcome Sanger Institute (https://wge.stemcell.sanger.ac.uk//find_crisprs). Annealed oligonucleotides containing the homology region were phosphorylated with T4 polynucleotide kinase (New England Biolabs, M201) and then ligated into the Bbs1 site of the vectors pX335 and pKN7 (Appendix Tables S1) in the presence of T4 DNA ligase (New England Biolabs, M202), as previously described (Pyzocha et al, 2014).

To create small deletions at a particular locus, mouse ES cells were transfected with two plasmids expressing the chosen pair of gRNAs together with the Cas9-D10A ‘nickase’ mutant and the puromycin resistance gene, as described below. *FAF1Δ UBXN7Δ* cells were produced by targeting the *UBXN7* locus in *FAF1Δ* cells. Similarly, *FAF1Δ UBXN7Δ FAF2-ΔUBX* cells were produced by targeting the *FAF2* locus in *FAF1Δ UBXN7Δ* cells.

### Transfection of mouse ES cells and selection of clones

A stock solution of 1 mg / ml linear Polyethylenimine (PEI; Polysciences, Inc 24765-2) was prepared in a buffer containing 25 mM Hepes, 140 mM NaCl, 1.5 mM Na_2_HPO_4_ adjusted to pH 7.0 with NaOH. The solution was sterilized by passing through a 0.2 µm filter and stored at −20 °C. 1 µg of each plasmid DNA (two plasmids expressing Cas9 and gRNAs) were mixed in 100 µl of reduced-serum medium OPTI-MEM (ThermoFisher Scientific, 31985062) then mixed with 15 µl of 1 mg/ml PEI, vortex well and left for 15 minutes. Subsequently, 1.0 × 10^6^ cells were aliquoted and centrifuged at 350 x g for 5 minutes before resuspension in 200 µl OPTI-MEM. The cell suspension and DNA-PEI mix were gently mixed and incubated at room temperature for 30 minutes in a 1.5 ml microfuge tube. Cells were then transferred to a single well of a 6-well plate that contained 2 ml of complete DMEM medium. Cells were incubated for 24 hours after transfection, followed by two 24-hour rounds of selection with fresh medium containing 2 µg / ml Puromycin (ThermoFisher Scientific, A1113802). The surviving cells were then released from the wells with 0.05 % Trypsin / EDTA, diluted 5 / 25 / 125 times with complete DMEM medium and then plated on 10 cm plates precoated with PBS containing 0.1 % (w/w) gelatin. Subsequently, colonies were expanded and monitored as appropriate by immunoblotting, PCR and DNA sequencing of the target locus.

### Genotyping of mouse ES cells by PCR and DNA sequencing

Cells from a single well of a 6-well plate were resuspended in 100 µl of 50 mM NaOH and the sample was then heated at 95 °C for 15 minutes. Subsequently, 11 µl of 1 M Tris-HCl (pH 6.8) was added to neutralise the pH. A 0.5 µl aliquot of the resulting genomic DNA solution was used as the template for genotyping purposes in 25 µl PCR reactions with Ex Taq DNA polymerase (TaKaRa, RR001). PCR products were subcloned with the TOPO TA cloning Kit (Invitrogen, K457540) and sequenced with T3 primer (5′-AATTAACCCTCACTAAAGGG-3′).

### Extracts of mouse ES cells and immunoprecipitation of protein complexes

1.2 x 10^7^ mouse ES cells were plated on a 15 cm petri dish and grown for 24 hours at 37 °C. Cells were released from the dishes by incubation for 10 minutes with 10 ml of PBS containing 1 mM EGTA, 1 mM EDTA and then harvested by centrifugation at 350 x g for 3 minutes, before snap-freezing with liquid nitrogen and storage at −80°C. Cell pellets from two 15 cm petri dishes were resuspended with an equal volume of ‘Cell lysis buffer’ supplemented with 5 μM Propargyl-Ubiquitin (MRC PPU Reagents and Services, DU49003) to inhibit deubiquitylase activity, and chromosomal DNA was then digested for 30 minutes at 4 °C with 1600 U / ml of Pierce Universal Nuclease (ThermoFisher Scientific, 88702). The extracts were centrifuged at 20,000 x g for 30 minutes at 4 °C. The supernatant was then mixed with 100 μl slurry of Protein A/G Sepharose beads (Expedeon, AGA1000) that had been pre-washed in Cell lysis buffer. This step was used to remove proteins that bound non-specifically to the beads. After 15 minutes of rotation at 4°C, the samples were centrifuged at 800 x g for 30 seconds at 4°C and the supernatant was recovered. A 20 μl aliquot of the supernatant was added to 40 μl of 1.5X LDS sample loading buffer.

The rest of the extract was then incubated for 90 minutes with 10 μl slurry of GFP-Trap Agarose beads (gta-100, Chromotek). The beads were washed four times with 1 ml of Wash buffer and bound proteins were eluted at 95 °C for 3 minutes in 30 μl of 1X LDS sample loading buffer.

### Inactivation of CUL2^LRR1^ in mouse ES cells

Lipofectamine™ RNAiMAX Transfection Reagent (ThermoFisher Scientific, 13778075) was used to introduce siRNA into mouse ES cells, according to the manufacturer’s protocol. *LRR1* siRNA (Horizon Discovery, J-057816-10) were transfected at a final concentration of 25 nM. After 24 hours, the transfection was repeated and incubation continued for a further 24 hours. Subsequently, cells were treated with 5 µM MLN-4924 (Activebiochem, A1139) for 5 hours before imaging. MLN-4924 inhibits the E1 enzyme of the protein neddylation pathway and thereby inhibits cullin ligase activity (Soucy et al., 2009).

### Imaging of mouse ES cells by spinning disk confocal microscopy

Confocal images of live cells were acquired with a Zeiss Cell Observer SD microscope with a Yokogawa CSU-X1 spinning disk, using a HAMAMATSU C13440 camera with a PECON incubator, a 60X 1.4-NA Plan-Apochromat oil immersion objective, and appropriate excitation and emission filter sets. Images were acquired using the ‘ZEN blue’ software (Zeiss) and processed with ImageJ software (National Institutes of Health) as previously described (Sonneville et al., 2017).

For time-lapse live cell imaging, mouse ES cells expressing GFP-SLD5 from the endogenous locus were grown on ‘µ-Dish 35 mm, high’ (Ibidi, 81156) with ‘no phenol red DMEM medium’ (ThermoFisher Scientific, 21063029) supplemented as described above. Imaging was started 48 hours after seeding, following treatment with siLRR1 and MLN-4924 as described above. Datasets were acquired every 6 minutes (2 × 2 binning, 0.25 seconds exposure for GFP), with heating to 37°C and 5% CO_2_.

### Cell survival assay for mouse ES cells

500 cells (or 1,000 cells for *FAF1Δ UBXN7Δ*) were plated into 10 cm petri dishes. On the next day, 0, 3, or 6 µl of 10 mM MLN-4924 or DMSO were added to 10 ml medium to give a final concentration of 0, 3, or 6 µM, before incubation for 6 hours. The drug-containing medium was then replaced with fresh medium, and incubation proceeded for a further six days. Cells were then washed with PBS and fixed with methanol. The fixed cells were stained with 0.5 % crystal violet solution (Sigma-Aldrich, HT90132), and images of the plates were captured with a scanner. The number of colonies was counted manually and the survival rate was calculated.

## Acknowledgements

We are grateful for the support of the Medical Research Council (core grant MC_UU_12016/13) and Cancer Research UK (Programme Grant C578/A24558). We thank Tom Deegan for help with reconstituted ubiquitylation reactions with budding yeast proteins, Cristian Polo Rivera for purified CMG, Yisui Xia for help with equivalent reactions involving *C. elegans* factors and Fabrizio Villa for assistance with mouse ES cell work. We are also grateful to Axel Knebel and Clare Johnson for purified Uba1 and Ubiquitin derivatives and MRC PPU Reagents and Services (https://mrcppureagents.dundee.ac.uk) for antibody production.

---

**Figure 3-figure supplement 1.**
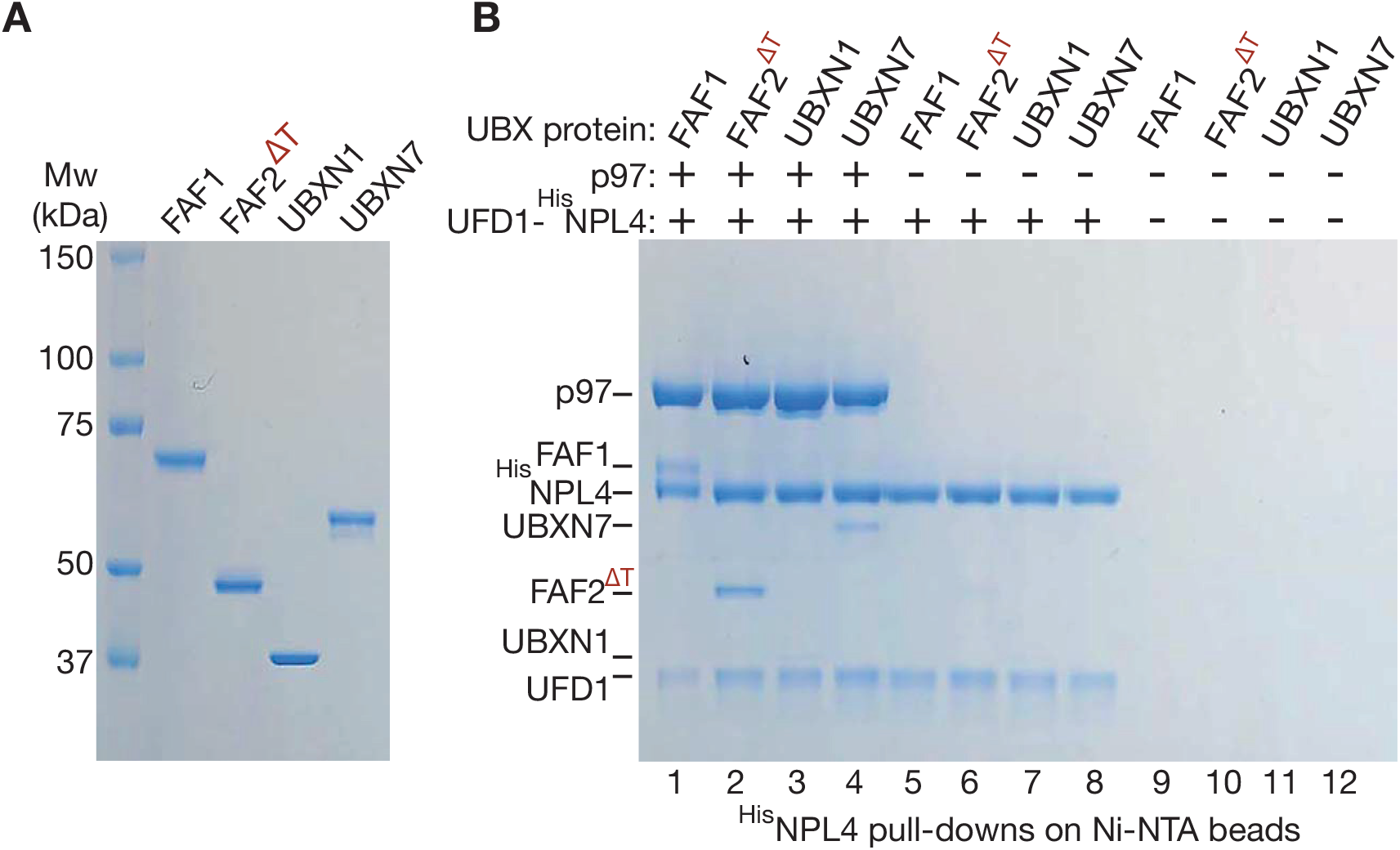
Purified human UBX proteins bind to p97-UFD1-NPL4. (**A**) Purified UBX proteins. (**B**) Proteins were mixed as indicated and the factors associated with ^His^NPL4 were isolated on Ni-NTA beads as described in Materials and Methods. The bound proteins were monitored by SDS-PAGE and Coomassie staining.

**Figure 3-figure supplement 2.**
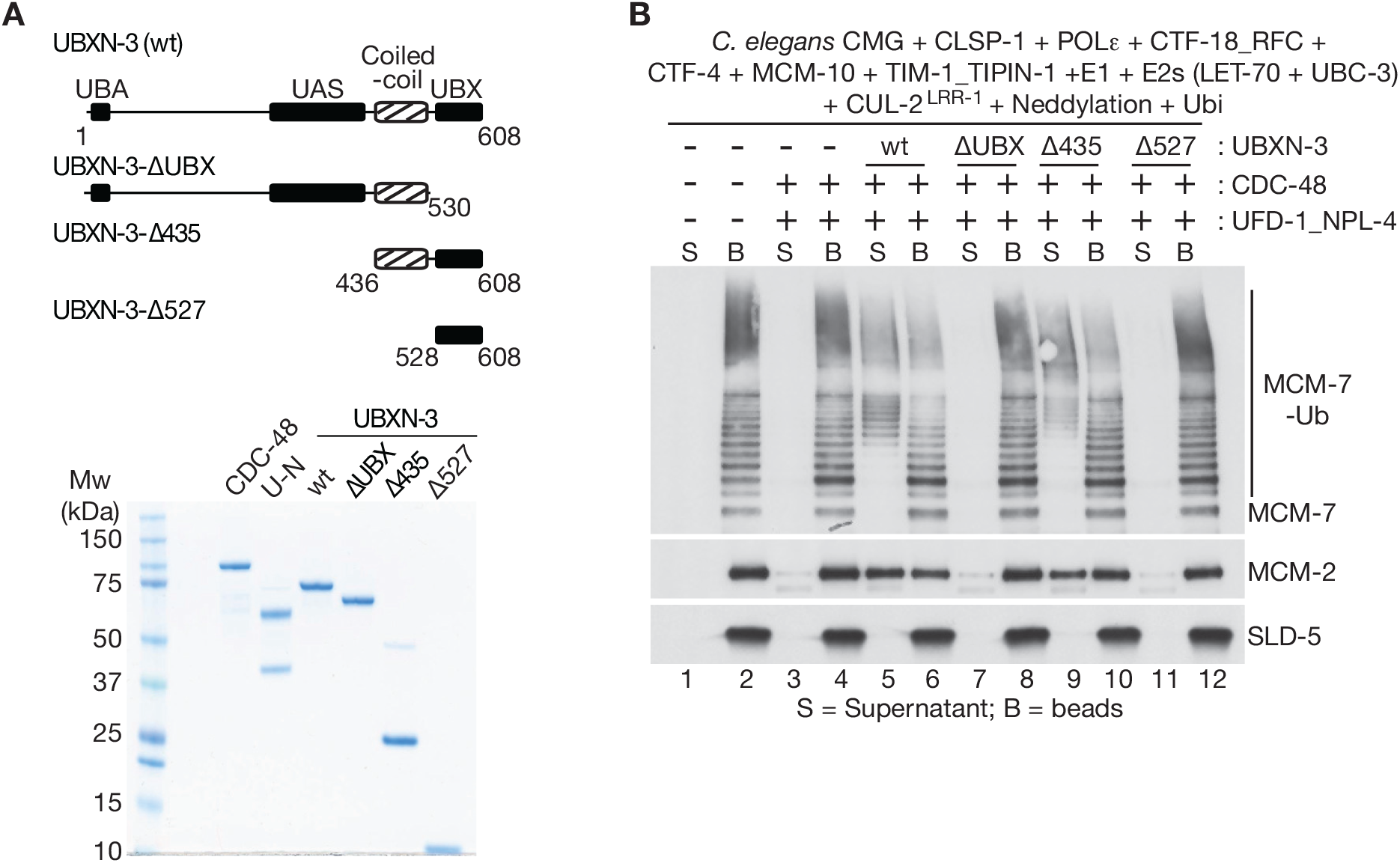
The coiled-coil and UBX domains of *C. elegans* UBXN-3 are required to stimulate the unfoldase activity of CDC-48_UFD-1_NPL-4. (**A**) The indicated truncations of *C. elegans* UBXN-3 (upper panel) were purified together with *C. elegans* CDC-48 and UFD-1_NPL-4 (lower panel; UN = UFD-1_NPL-4). (**B**) *C. elegans* CMG was ubiquitylated in the presence of the indicated factors and then bound to beads coated with antibodies to the SLD-5 subunit of GINS, as described in Materials and Methods. Ubiquitylated CMG was then incubated as shown with *C. elegans* CDC-48, UFD-1_NPL-4 and wt or truncated forms of UBXN-3. CMG disassembly was monitored by release of ubiquitylated MCM-7 (MCM-7-Ub) and MCM-2 into the supernatant.

**Figure 5-figure supplement 1.**
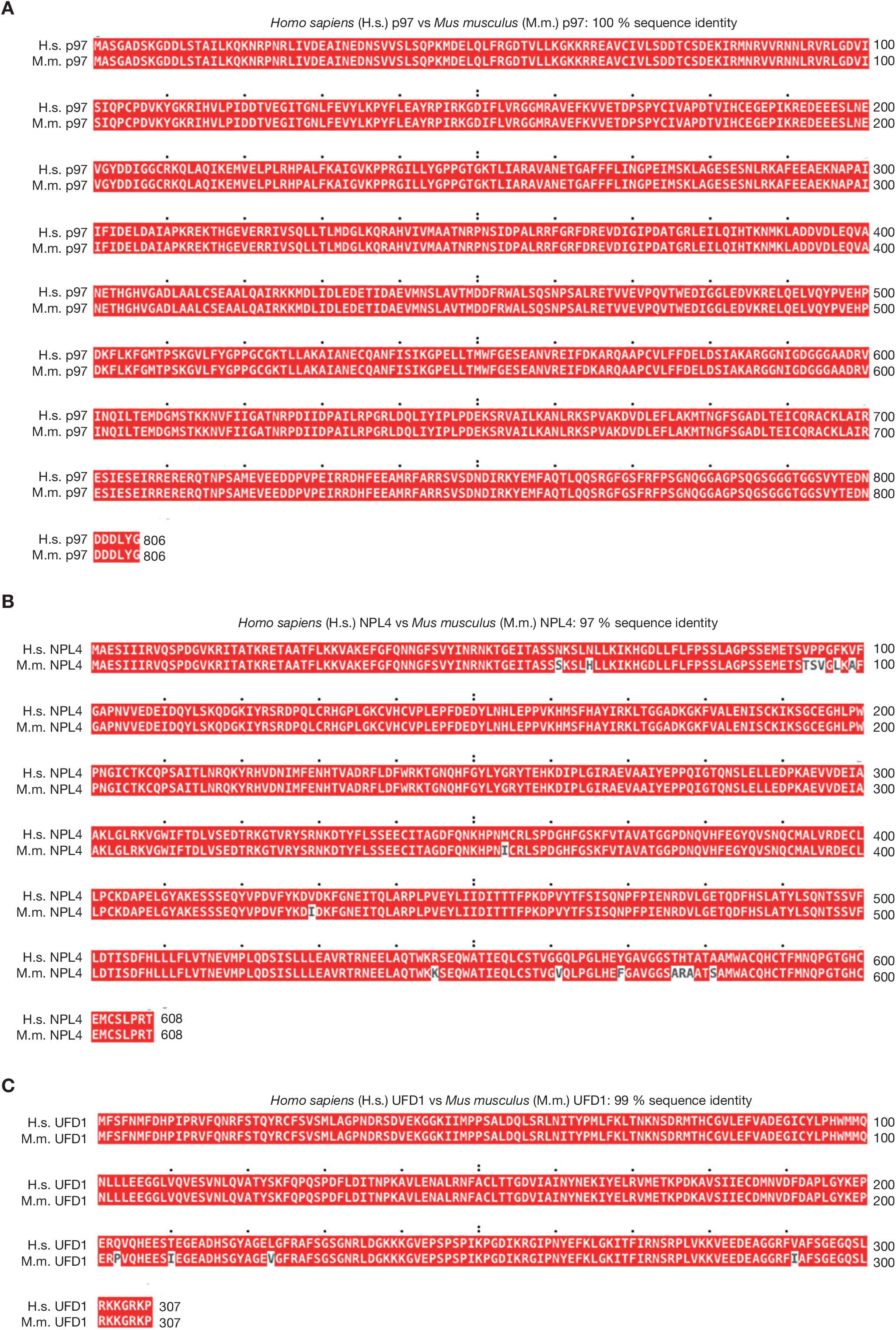
Sequence alignment of human and mouse orthologues of p97 and UFD1-NPL4. (**A**) Human and mouse orthologues of p97 were aligned with Clustal Omega software and the output viewed with MView (Madeira et al., 2019). (**B**-**C**) Similar analysis for NPL4 and UFD1.

**Figure 5-figure supplement 2.**
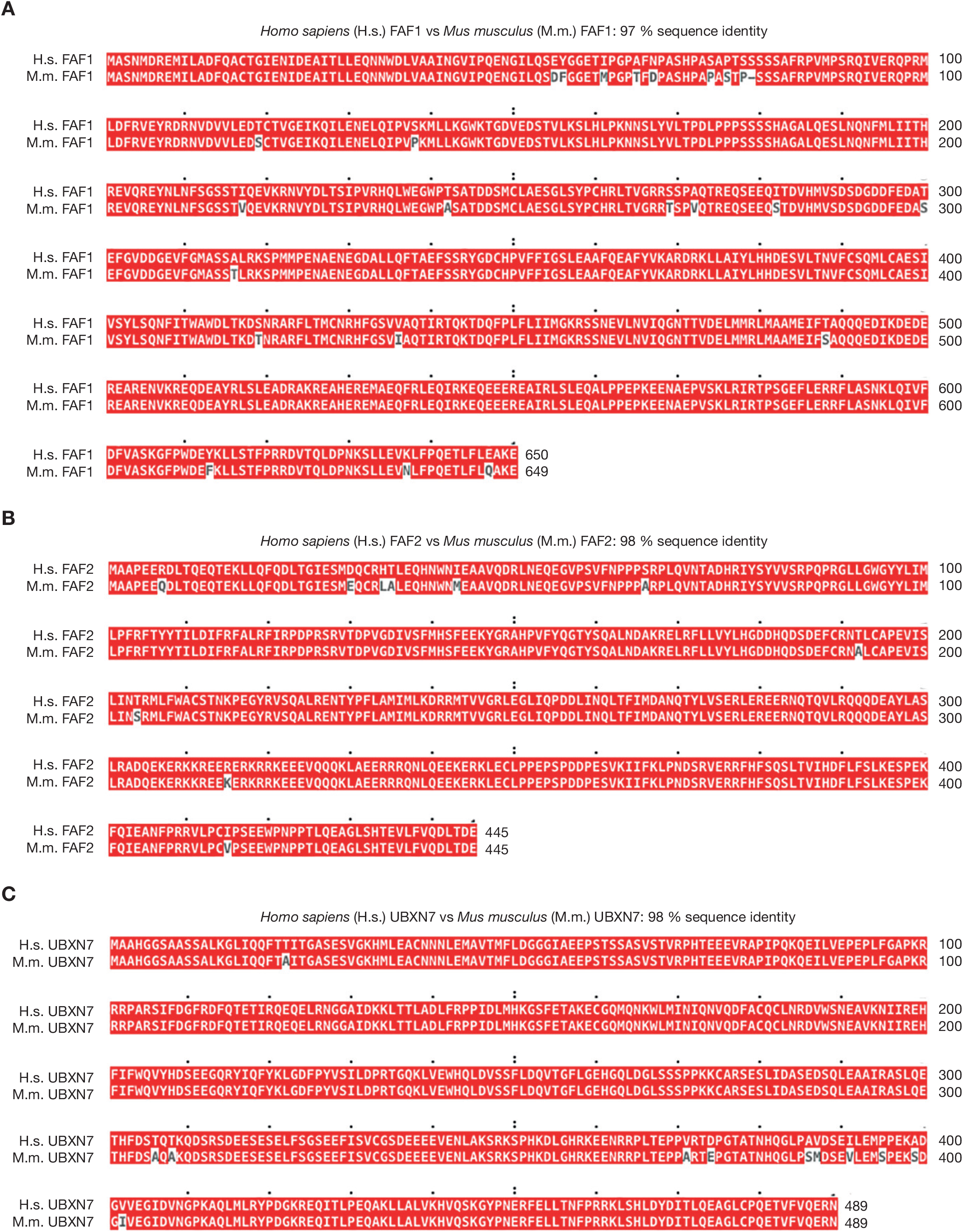
Sequence alignment of human and mouse orthologues of FAF1, FAF2 and UBXN7. (**A**) Human and mouse orthologues of FAF1 were aligned with Clustal Omega software and the output viewed with MView (Madeira et al., 2019). (**B**-**C**) Similar analysis for FAF2 and UBXN7.

**Figure 5-figure supplement 3.**
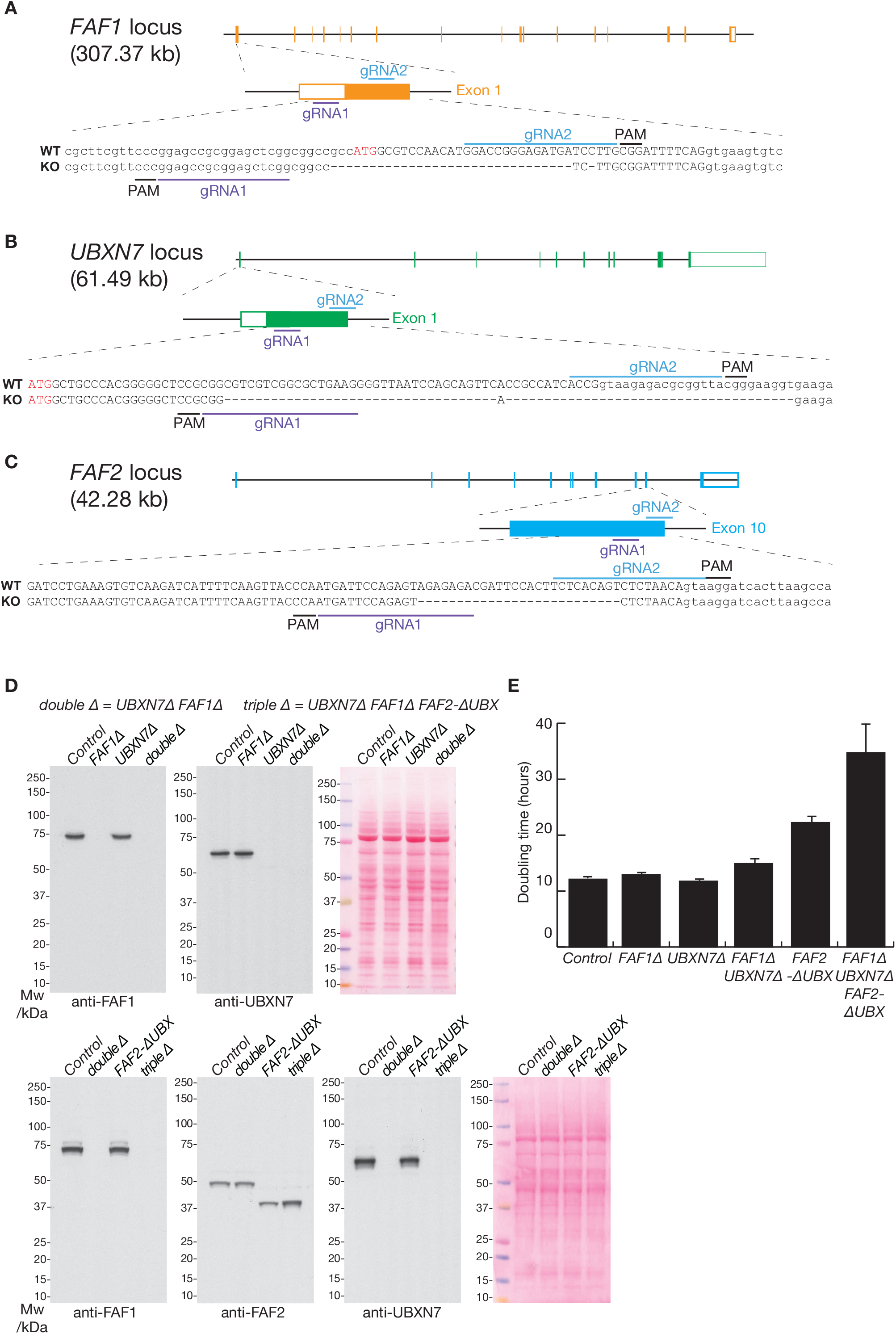
Deletion of the *FAF1*, *UBXN7* and *FAF2* genes by CRISPR-Cas9. (**A**-**C**) Loci and corresponding guide RNAs (gRNAs) that were used to target the D10A ‘nickase’ mutant of Cas9 to the *FAF1*, *UBXN7* and *FAF2* loci in mouse ES cells. See Materials and Methods for further details. PAM = ‘Protospacer Adjacent Motif’ that takes the form ‘NGG’ and is essential for cutting by Cas9. (**D**) Immunoblot analysis with the indicated antibodies to monitor deletion of the *FAF1*, *FAF2* and *UBNX7* genes. The right panels show total protein stained with Ponceau S. (**E**) Doubling time of mES cells with the indicated genotypes was monitored as described in Materials and Methods. The histogram indicates the mean values from three biological replicates, together with the associated standard deviations.

**Figure 5-figure supplement 4.**
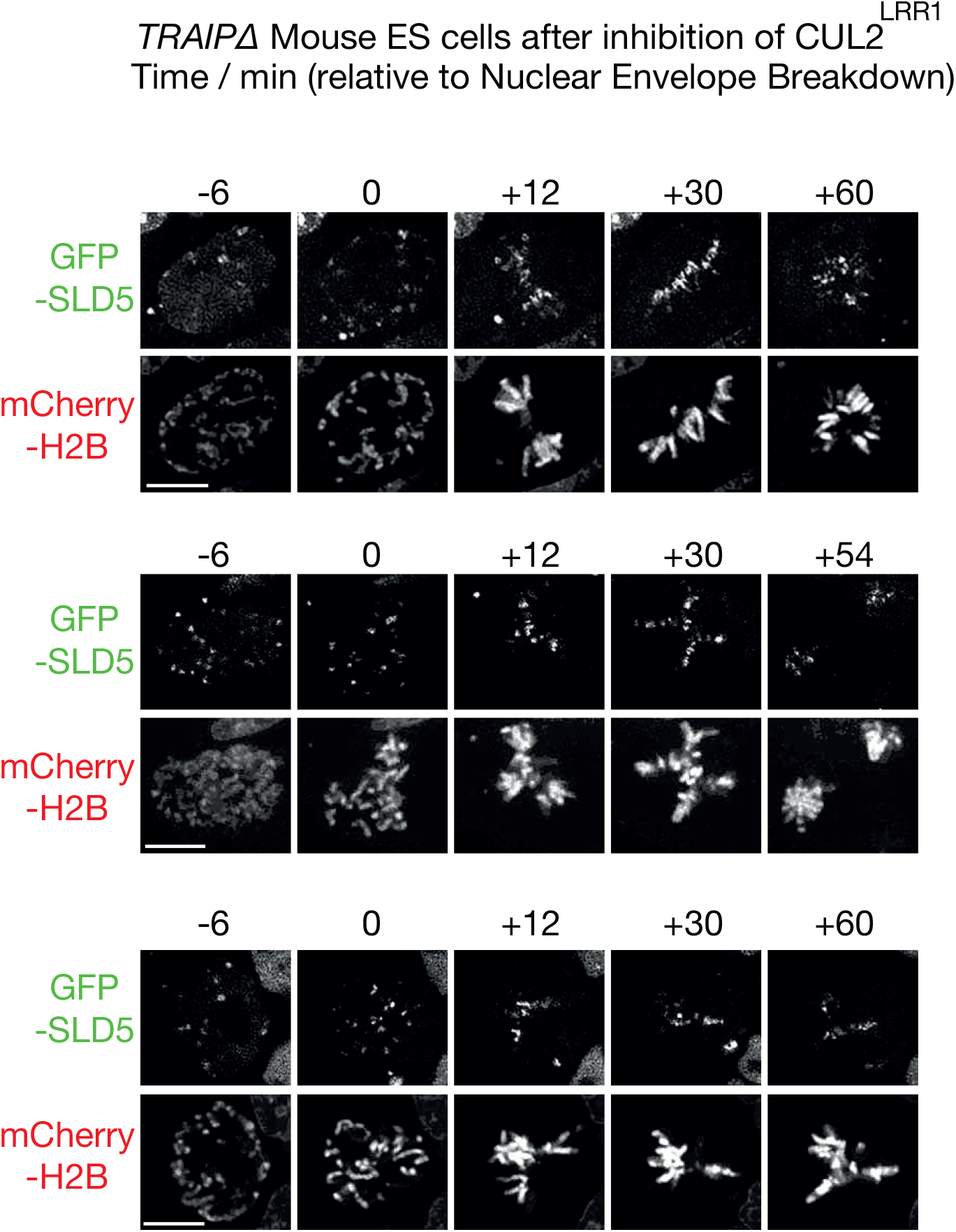
Mitotic CMG disassembly is blocked in mES cells that lack TRAIP. TRAIPΔ cells expressing *GFP-SLD5* from the endogenous *SLD5* locus were grown as in Figure 5C and monitored by time-lapse video analysis. Three examples are shown of mitotic entry following inhibition of CUL2^LRR1^, illustrating the persistence of GFP-SLD5 on mitotic chromatin throughout the observed time course (100 % of cells, n = >20 cells per sample). Scalebars correspond to 10 µm.

